# Are isokinetic leg torques and kick velocity reliable predictors for competitive success in taekwondo athletes?

**DOI:** 10.1101/2020.06.19.161158

**Authors:** Pedro Vieira Sarmet Moreira, Coral Falco, Luciano Luporini Menegaldo, Márcio Fagundes Goethel, Leandro Vinhas de Paula, Mauro Gonçalves

**Affiliations:** Biomedical Engineering Program – COPPE – Federal University of Rio de Janeiro- RJ, Brazil; Laboratory of Biomechanics, State University of São Paulo (UNESP), Rio Claro-SP, Brazil; Department of Sport, Food and Natural Sciences, Western Norway University of Applied Sciences, Bergen, Norway; Laboratory of Biomechanics, Federal University of Minas Gerais, Belo Horizonte-MG, Brazil

**Keywords:** Isokinetic torque, kinematic analysis, kick performance, velocity, impact, discriminant analysis

## Abstract

The aim of the study was to analyzed the relationship between isokinetic knee and hip peak torques and Roundhouse-kick velocities and expertise level (Elite vs. Subelite) of Taekwondo athletes. Seven elite and seven sub-elite athletes were tested for kick kinematic, power of impact and for isokinetic peak torque (PT) at slow (60°/s) and high (240°/s) concentric mode. PTs were compared between groups and correlated with the data of kick performance. It was found inter-group differences in hip flexors and extensors PT at the isokinetic fast speed. The hip flexion PT at 60°/s and 240°/s were negatively correlated with the kick time (R = −0.46, and R = −0.62, respectively). Hip flexion torque at 60°/s was also positively correlated (R = 0.52) with the peak of linear velocity of the foot (LVF) and the power of impact (R = 0.51). Peak torque of hip extension at 60°/s and hip abduction at 240°/s were correlated with the LVF (R= 0.56 and R = 0.46). Discriminant analysis presented an accuracy of 85.7% in predicting expertise level based on fast torques of hip flexion and extension and on the knee extension velocity during the kick. This study demonstrated that hip muscles strength is probably the dominant muscular factor for determining kick performance. Knee angular velocity combined with hip torques are the best discriminators for the competitive level in taekwondo athletes.

## Introduction

The most popular technique in Taekwondo (TKD) combats is the Roundhouse Kick, or Bandal Chagui [1-3]. Defined as a multiplanar and multi-joint action[2-6], is described as a *proximo-distal sequence*, wherein structures nearest the center of the body (proximal segments) develop first in temporal order of joints movement, while distal segments lag behind, and are thereafter followed by its relative distal segment acceleration while proximal segments decelerate [7-9].

According to this principle, the largest possible velocity of the proximal segment, linked by the interaction with the following segments, determines the terminal velocity or impact magnitude [3-5,7,9-13]. Angular acceleration of a segment is caused by muscle torques that control the proximal joint [6,7,15] and by the angular momentum transmitted to the next (distal) segment [7,15,16]. The resultant torque produced during the kick depends on the athlete’s coordinative capacity to maximize the agonist and minimize the antagonist torque [7,17,18].

During a kick, torques are generated at either low or high angular velocities [7,19], which can be reasonably reproduced by isokinetic dynamometers [20-22]. Such instruments present high reliability [23], validity [22], and control of speed and range of motion [22,23]. Previous researchers [6,24] demonstrated that elite athletes of combat sports presented improved capacities to produce torque compared with sub-elite athletes or non-athletes, in isokinetic velocities from 30°/s to 400°/s. Fong & Tsang [21] considered the velocity of isokinetic contractions (by using an isokinetic device) of 60°/s as low and 240°/s as high speed. The device controls limit the capacity to produce the movement but not the capacity of the athletes to produce the most forceful and explosive contractions. In this sense, the contraction produced would be related to the torque along time or to the angular momentum (inertial moment multiplied by the angular velocity). Authors showed that the number of TKD training hours per week positively correlated with knee extensors (R = 0.639) and flexors’ (R = 0.472) peak torque at 240°/s, but not at 60 °/s, nor with the ankle plantar flexors at any speed of contraction. Pieter et al. [25] showed that TKD practitioners reach higher hamstring isokinetic torques than non-athletes, at different velocities of contraction. Elite TKD athletes have better performance than sub-elite athletes, kicking faster and stronger, and presenting higher angular and linear velocities, and lower kick times [5,26,27]. However, the isokinetic torque of TKD athletes has not yet been associated with the specific performance of martial kicks.

Moreover, it can be difficult to know if an athlete has the physical and technical performance corresponding to the biomechanical performance of an athlete of elite or sub-elite level. In this sense, the analysis of specific biomechanical parameters can help us to rank and characterize them, built on a discriminant analysis (DA). DA is a predictive method to characterize two or more classes of people based on means of classifiers for dimensionality reduction before later classification. DA is used when groups are a priori known. Each subject (case) must have a score in one or more quantitative predictor measures, and a score on a group measure [28] Discriminant function analysis is a classification. The predictive model of a group membership according to the athlete’s level can be useful to rank athletes based on the selected parameters.

Thus, three hypotheses will be addressed: 1) elite athletes show higher isokinetic torques than sub-elite athletes, regardless of the muscle group and contraction speed; 2) isokinetic torques correlate positively with peak linear and angular velocities and impact magnitudes obtained during the kick, and negatively with the temporal parameters of the kicks and; 3) a discriminant equation variables is a strong predictor for the ranking performance.

## Material and methods

### Participants

Fourteen black-belt TKD athletes, divided into two groups: 7 elite athletes (that at least obtained a medal in a national Brazilian championship: 23.6 ± 2.1 years; 69 ± 9.5 kg; 168 ± 5 cm; 12.2 ± 8.5 years of training; 15.7 ± 4.7 hours per week of training), and 7 sub-elite athletes (whose best result was a participation in a national competition: 22.4 ± 1.3 years; 66.8 ± 14.2 kg; 174 ± 11 cm; 10.4 ± 6.1 years of training; 11.4 ± 5.1 hours per week of training), participated in the study. Preliminary analysis showed no significant differences between groups (p > 0.05), in any of the variables. The study was approved by the ethic committee (n. 058/2013), and all participants signed an informed consent form.

### Experimental design

Participants were first evaluated through kinematic analysis and impact measurement during the kick execution. After 10 minutes of passive rest, knee and hip isokinetic torque curves were measured at two different velocities (60°/s and 240°/s).

### Data collection

Kinematics were measured using seven MX13 cameras (Vicon®) sampled at 250 Hz. Thirty-nine marker reflectors were placed on each athlete, according to the “*Plugin Gait Full Body (UPA and FRM)*” (Vicon®) marker set. After 15 minutes of warm-up, 9 Bandal Chagui kicks were performed directed to a dummy (=BoomBoxe®, see video 1: https://doi.org/10.6084/m9.figshare.9698741.v2), instrumented with a trunk protector (TK-Strike 4.2, Daedo®) used for official competitions. The trunk protector registered the impact power in a measurement unity appropriate for the WT, but not defined by the International System of Units. A force plate OR-6 (AMTI®) sampled at 2000 Hz, placed on the rear dominant kicking leg, determined the kick start onset.

The isokinetic protocol consisted of a three submaximal and one maximal familiarization contractions, followed by 15 seconds of rest and five maximal contractions, alternating between reciprocal movements of knee flexion/extension, hip flexion/extension, and hip adduction/abduction at 60°/s e 240°/s speeds. Between the pairs of reciprocal movements, and during the alternation of speed contraction, there were two minutes of rest intervals. The order of the velocities and the pairs of movement were randomized. Participants were instructed to make the contractions “as fast and strong as possible”. There was verbal encouragement during the contractions. Torque-angle curves were collected, using an isokinetic dynamometer System 4 PRO (Biodex^®^), at a sampling frequency of 3000Hz by, an A/D converter module (Noraxon^®^, NorBNC) and MR version 3.2 (Noraxon^®^) software.

Participants stayed seated with the chair backrest fixed by straps to the chair, adjusted in 110°from the horizontal surface. The dynamometer axis aligned with the greater trochanter of the femur. The knee started at 90°flexion, extended 90^o,^ and returned to the initial position. During the hip evaluation in the sagittal plane, the thigh started from 5°of flexion (0°= full extension) and covered 50°of flexion/extension. For the hip abduction/adduction evaluation, participants were positioned in the lateral decubitus and the knee fully extended, with the rotational dynamometer axis aligned with the Ischiatic Tuberosity and the dynamometer level fixed to the dominant thigh. The range of hip abduction/adduction was between 0°and 45°, with 0°defined by the legs at the horizontal position.

### Data Processing

The highest peaks of isokinetic torques, linear and angular peak velocities of the hip, knee, and ankle, as well as impact power, were selected for further analysis. Kinematics and ground reaction forces (GRF) were smoothed through a 4^th^ order low-pass *Butterworth zero-lag* filter, with cut-off frequencies of 85Hz and 10Hz, respectively. Three kick events (onsets) were identified: *t*_*1*_ - preparation time; *t*_*2*_ - kicking time; and *t*_*3*_ - impact time. *t*_*1*_ is the time instant corresponding to an initial 2.5% variation of the GRF with respect to the basal value. *t*_*2*_ is time instant corresponding to when the GRF turns zero. *t*_*3*_ is when the foot touches the target, i.e., when the dummy contact sensor overcomes a voltage threshold. Next, the preparation time (PT= *t*_*2*_ – *t*_*1*_) and kicking time (KT *t*_*3*_ – *t*_*2*_) were calculated. They correspond to the time while the foot performs impulse against the ground (PT) and performs the aerial trajectory from the ground to the target (KT). During the kicking time, the maximal value obtained in the velocity- curve, each linear and angular movement were used for determining the linear (Pelvis: Anterior Superior Iliac Crest; Knee: Lateral Condyle of Femur; and Foot: Center of Gravity of the Foot) and angular peak velocities (Hip: Flexion, Extension, Adduction and Abduction; Knee: Flexion and Extension) of the leg anatomical points. The collected torque vectors were smoothed with a 4^th^ order low-pass *Butterworth zero-lag* filter with a cut-off frequency of 20 Hz and corrected for segment weight according to equation 1.

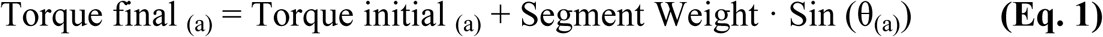

where “a” represents the sample and “θ” the segment angle of the vertical orientation.

### Statistical Analysis

Normality of the data was verified through the *Shapiro-Wilk* and homogeneity by *Levene* tests. Analysis of Variance (ANOVA) or a Mann-Whitney test, for parametric and no parametric statistics, respectively, was used to compare isokinetic torque, impact magnitude and selected kinematic data of kick speed, between groups. *Pearson* or *Spearman* correlation, for parametric and non-parametric statistics, respectively, was used to analyze the relationship between the kinematics and the impact magnitudes obtained during the kick. *Cohen’s f* scores [29], was used to quantify the effect size of the comparisons. An *f* value of 0.4, 0.25 and 0.1 was considered large, moderate and small effect, respectively.

DA was performed for the isokinetic torques and kinematic data to investigate how the general muscle torque and specific kick performance can discriminate the expertise level: sub-elite = 1; elite = 2. Only data that presented linearity, normality, multi-collinearity, homogeneity of variances, multivariate-normal distribution, and significant difference between groups through ANOVA were used in a stepwise discriminant analysis with the expertise level. Only two torque variables (hip flexion and hip extension, both at 240 °/s) and two kinematic variables (AVKnExt: angular velocity of knee extension and LFV: linear velocity of the knee) met all the necessary assumptions to perform the DA.

The first DA generated a general equation to classify the athletes based on the input variables. Thereafter, the cross-validation was performed based on the method Lachenbruch *U*. That is, the successive classifications of all cases but one to develop a discriminant function and then, categorize the case that was left out. The process was repeated with each case left out in turn and produces a more reliable function. Lastly, a “Matrix of Classification” was generated, containing the number of correct and incorrect athletes classified. Main metrics to classify the performance, based on the cross validated Matrix of Classification, was calculated based on if athletes were correctly classified, into four alternatives: True Elite (TE), False Elite (FE), True Sub-elite (TS) and False Sub-elite (FS). Three main metrics of performance were identified:

1. Sensitivity: the percentage of actual elite athletes correctly identified as elite, according to the equation 2.

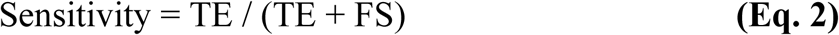
2. Specificity: the percentage of sub-elite athletes correctly identified as sub-elite, according to the equation 3:

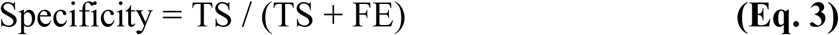
3. Accuracy: the percentile of participants correctly classified in the overall data, according to the equation 4:

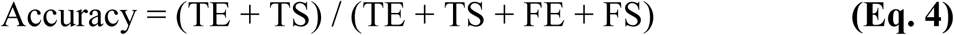

The effect sizes were calculated with the *GPower*® 3.1 (Dusserdolf, GE) software, using the method described by Faul et al.,[30,31]. The other statistical tests were performed with the software SPSS 18.0 (SPSS Inc., Chicago, IL) considering *p* < 0.05 as the significance level.

## Results

Table 1 shows comparative data between groups of kinematic and dynamic variables obtained during the kick and torques during the isokinetic evaluations. A Mann Whitney test indicated that kicking time was significantly lower for the elite than for the sub-elite group (U = 11.0, p = 0.042). During the kick, peak angular velocity during knee flexion was significantly higher for the elite than for the sub-elite group (U = 11.0, p = 0.042). ANOVA indicated that the linear peak velocity of the knee marker (F_(1,12)_ = 4.22, p = 0.031) and the angular velocity of knee extension (F_(1,12)_ = 6.18, p = 0.014) were higher in the elite than in the sub-elite group. Regarding isokinetic at 60 **°**/s and 240 **°**/s, from each joint, the results showed significant differences during hip flexion (F_(1,12)_ = 3.561, p = 0.042) and extension (F_(1,12)_ = 3.953, p = 0.035) in peak torque at 240 **°**/s between elite and sub-elite taekwondo athletes. Those differences were considered large (Cohen’s *f* > 0.50).

**Table 1.** Comparative data between groups of kinematic and dynamic variables obtained during the Bandal Chagui and torques during the isokinetic evaluations. **Legend: PT**, Preparation Time, the impulse phase of the kick; **KT**, Kick Time, the aerial phase of the kick; **LVF**, Linear Foot Velocity; **LVK**, Linear Knee Velocity; **LVP**, Linear Pelvis Velocity; **KnFlx**, Knee Flexion; **KnExt**, Knee Extension; **HipFlx**, Hip Flexion; **HipExt**, Hip Extension; **HipAdd**, Hip Adduction; **HipAbd**, Hip Abduction; **WTU**, World Taekwondo Federation (WT) patented Unity (U) of Power Impact. **°/s**, Isokinetic Velocity in Degrees for Second; **f:** Effect Size “f” from ANOVA test; ***η2***, percentage of explained variance of ranking classification in elite or sub-elite level; **\***, P < 0.05.

| Kind of Variable | Kinematic Variables | Athlete Groups |  | Effect Size |  |
| --- | --- | --- | --- | --- | --- |
| | | Elite | Sub-elite | (Cohen's *f*) | ($\eta^2$) |
| Time<br>(ms) | PT | 244±73 | 320±104 | 0.42 | 0.15 |
|  | **KT** | 213±9 | 237±36 | 0.46 | **0.17\*** |
| Peak Linear<br>Velocities<br>(m.s<sup>-1</sup>) | LVF | 17.36±0.70 | 16.53±1.35 | 0.38 | 0.13 |
|  | **LVK** | 8.20±0.44 | 7.73±0.41 | 0.55 | **0.23\*** |
|  | LVP | 3.14±0.34 | 3.03±0.28 | 0.17 | 0.03 |
| Peak Angular<br>Velocities (°/s) | **KnFlx** | 994±242 | 832±322 | 0.28 | **0.07\*** |
|  | **KnExt** | 1584±78 | 1406±173 | 0.66 | **0.31\*** |
|  | HipFlx | 451±96 | 428±196 | 0.07 | 0.01 |
|  | HipExt | 457±196 | 365±110 | 0.29 | 0.08 |
|  | HipAbd | 441±127 | 365±101 | 0.33 | 0.10 |
| Dynamic (WTU) | Impact | 44.57±1.33 | 44.66±2.20 | 0.01 | 0.00 |
| Isokinetic<br>Moment<br>(Nm*Kg-1) | KnFlx 60°/s | 1.60±0.18 | 1.60±0.24 | 0.00 | 0.00 |
|  | KnFlx 240°/s | 1.23±0.19 | 1.22±0.12 | 0.01 | 0.00 |
|  | KnExt 60°/s | 2.79±0.30 | 2.61±0.43 | 0.25 | 0.06 |
|  | KnExt 240°/s | 1.64±0.17 | 1.67±0.26 | 0.05 | 0.003 |
|  | HipFlx 60°/s | 2.22±0.31 | 2.02±0.32 | 0.33 | 0.10 |
|  | **HipFlx 240°/s** | 1.78±0.17 | 1.59±0.21 | 0.50 | **0.20\*** |
|  | HipExt 60°/s | 2.65± 0.41 | 2.36± 0.31 | 0.40 | 0.14 |
|  | **HipExt 240°/s** | 1.73±0.62 | 1.10 ± 0.56 | 0.53 | **0.22\*** |
|  | HipAdd 60°/s | 1.69±0.43 | 1.75± 0.41 | 0.07 | 0.01 |
|  | HipAdd 240°/s | 0.92±0.23 | 0.86±0.48 | 0.08 | 0.01 |
|  | HipAbd 60°/s | 1.93±0.36 | 1.83± 0.21 | 0.17 | 0.03 |
|  | HipAbd 240°/s | 1.13±0.43 | 1.06±0.23 | 0.10 | 0.01 |

Table 2 shows the correlations between the isokinetic peak torque and the roundhouse kick parameters.

**Table 2.**
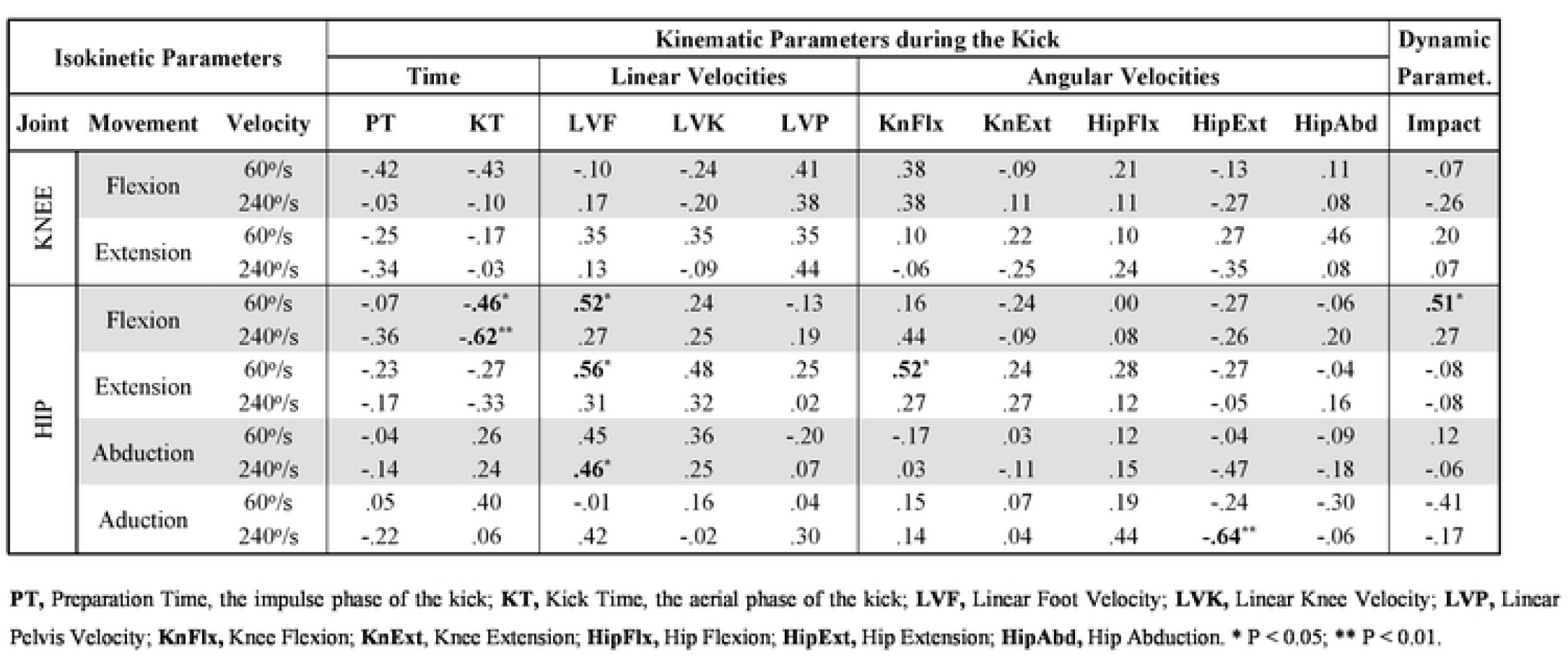
Correlations coefficients obtained between the isokinetic peak torques and the temporal, the kinematic and the impact data obtained during the Bandal Chagui kick (n=14 athletes as a unique group). **Legend: PT**, Preparation Time, the impulse phase of the kick; **KT**, Kick Time, the aerial phase of the kick; **LVF**, Linear Foot Velocity; **LVK**, Linear Knee Velocity; **LVP**, Linear Pelvis Velocity; **KnFlx**, Knee Flexion; **KnExt**, Knee Extension; **HipFlx**, Hip Flexion; **HipExt**, Hip Extension; **HipAbd**, Hip Abduction. * P < 0.05; ** P < 0.01.

| Isokinetic Parameters |  |  | Kinematic Parameters during the Kick |  |  |  |  |  |  |  |  |  | Dynamic Paramet. |
| --- | --- | --- | --- | --- | --- | --- | --- | --- | --- | --- | --- | --- | --- |
|  |  |  | Time |  | Linear Velocities |  |  | Angular Velocities |  |  |  |  |  |
| Joint | Movement | Velocity | PT | KT | LVF | LVK | LVP | KnFlx | KnExt | HipFlx | HipExt | HipAbd | Impact |
| KNEE | Flexion | 60°/s | -.42 | -.43 | -.10 | -.24 | .41 | .38 | -.09 | .21 | -.13 | .11 | -.07 |
|  |  | 240°/s | -.03 | -.10 | .17 | -.20 | .38 | .38 | .11 | .11 | -.27 | .08 | -.26 |
|  | Extension | 60°/s | -.25 | -.17 | .35 | .35 | .35 | .10 | .22 | .10 | .27 | .46 | .20 |
|  |  | 240°/s | -.34 | -.03 | .13 | -.09 | .44 | -.06 | -.25 | .24 | -.35 | .08 | .07 |
| HIP | Flexion | 60°/s | -.07 | -.46* | .52* | .24 | -.13 | .16 | -.24 | .00 | -.27 | -.06 | .51* |
|  |  | 240°/s | -.36 | -.62** | .27 | .25 | .19 | .44 | -.09 | .08 | -.26 | .20 | .27 |
|  | Extension | 60°/s | -.23 | -.27 | .56* | .48 | .25 | .52* | .24 | .28 | -.27 | -.04 | -.08 |
|  |  | 240°/s | -.17 | -.33 | .31 | .32 | .02 | .27 | .27 | .12 | -.05 | .16 | -.08 |
|  | Abduction | 60°/s | -.04 | .26 | .45 | .36 | -.20 | -.17 | .03 | .12 | -.04 | -.09 | .12 |
|  |  | 240°/s | -.14 | .24 | .46* | .25 | .07 | .03 | -.11 | .15 | -.47 | -.18 | -.06 |
|  | Aduction | 60°/s | .05 | .40 | -.01 | .16 | .04 | .15 | .07 | .19 | -.24 | -.30 | -.41 |
|  |  | 240°/s | -.22 | .06 | .42 | -.02 | .30 | .14 | .04 | .44 | -.64** | -.06 | -.17 |

Regarding the DA analysis, the results showed that (table 3), in the first analysis, hip flexion (TqHF_240_) and extension (TqHE_240_), at 240°/s, predicted competitive level (multiple correlation with the discriminant equation: R = 0.61 and R = 0.58, respectively). The discriminant function revealed a significant association between the groups and all predictors, accounting for 47% of the between-groups variability. The cross-validated classification showed by the accuracy that 64.3% of the participants were correctly classified.

**Table 3.** Discriminant Analysis Result of the Discriminant Functions (D_1,_ D_2_ and D_3)_. **D**_**1**_: Discriminant equation based on isokinetic torque data; **D**_**2**_: Discriminant function based on kinematic data of kick performance; **D**_**3**_: Discriminant equation based on isokinetic torques and kinematic data of kick performance; **AvKnExt:** Peak of Angular Velocity of Knee Extension; **LKV:** Peak of Linear Velocity of the Knee Marker; **TqHE**_**240**_: Peak Torque of Hip Extension at 240°/s;**TqHF**_**240**_: Peak Torque of Hip Flexion at 240°/s; **Sensitivity:** the percentage of actual positives (Elite) that are correctly identified as positive; **Specificity:** measures the percentage of failures (Subelite) correctly identified as failures; **Accuracy:** the percentage of athletes correctly classified in their correspondent group (Elite or Subelite); All the presented equations and metrics are based on the Cross Validated Matrix of Classification.

**Figure 3 – Discriminant Analysis Result of the Discriminant Functions (D<sub>1</sub>, D<sub>2</sub> and D<sub>3</sub>).**
| Discriminant Equations |  |  |  |  | Centroid Predictive Value |  | Canonical Correlation |  |
| --- | --- | --- | --- | --- | --- | --- | --- | --- |
|  |  |  |  |  | Elite | Subelite | R | p |
| **D1** = TqHE<sub>240</sub> * 1.454 + TqHF<sub>240</sub>* 4.337 - 9.352 |  |  |  |  | 0.872 | -0.872 | 0.686 | 0.003 |
| **D2** = AVKnExt * 0.0074 - 11.13 |  |  |  |  | 0.665 | -0.665 | 0.583 | 0.029 |
| **D3** = AVKnExt * 0.0074 + TqHE<sub>240</sub> * 1.154 + TqHF<sub>240</sub> * 5.44 -21.8 |  |  |  |  | 1.542 | -1.542 | 0.857 | 0.003 |
| Equations | TE (%) | FE (%) | TS (%) | FS (%) | Sensitivity (%) |  | Specificity (%) | Accuracy (%) |
| **D1** | 4 (57.1) | 2 (28.6) | 5 (71.4) | 3 (42.9) | 57.1 |  | 71.4 | 64.3 |
| **D2** | 6 (85.7) | 2 (28.6) | 5 (71.4) | 1 (14.3) | 85.7 |  | 71.4 | 78.6 |
| **D3** | 6 (85.7) | 1 (14.3) | 6 (85.7) | 1 (14.3) | 85.7 |  | 85.7 | 85.7 |

In the second analysis, the predictors were kinematic variables (AVKnEx and LFV). The discriminant function revealed a significant association between groups and only one significant predictor for the model (AVKnExt), accounting for 34% of the between- group variability. The cross-validated classification presented an accuracy of 78.6%. Finally, a third analysis combined both, the peak torque and kinematical significant variables of the previous DA. The discriminant function revealed a significant association between the groups and all the predictors, accounting for 74% of the between-groups variability. The structure matrix revealed that all the variables were significant for the model, namely AVKnExt (R = 0.43), TqHE_240_ (R = 0.34) and TqHF_240_ (R = 0.33). The cross-validated classification showed an accuracy of 85.7%.

## Discussion

The present study aimed to compare two groups (elite and sub-elite) of taekwondo athletes and evaluate whether the peak isokinetic torques during the hip and knee (flexion, extension, abduction and adduction), at two different velocities of contraction (60 °/s and 240 °/s), were associated with the kinematic and dynamic parameters during a roundhouse kick. Also, based on discriminant analysis, build an equation to rank athletes’ performance predicted by relevant variables from the kinematic and peak torque data.

Regarding the first hypothesis, elite and sub-elite athletes presented significant differences in their peak isokinetic torque values, in favor of the elite group. These differences were significant in the hip sagittal plane (flexion and extension), at a high velocity (240°/s). As the athletes did not differ in volume and frequency of training, the difference can be due to, at least, one of two factors: 1) training quality and 2) conditional genetic factors for muscle power production, since differences were found only in the high speed of contraction. We find the first as the most probable explanation since the differences between groups manifested themselves only for specific muscle groups. If this is the case, we can suggest that the quality of force training with high speed of movements for sagittal hip movements should have a significant effect on the sport-specific classification.

When it comes to the velocity of contraction, high and low isokinetic torque velocity during the hip movement correlated with essential parameters (namely KT, LFV, KnFlex, angular velocity and impact) of the kinematical performance of the kick. It is possible to explain the significant correlations for the fast isokinetic contraction by the principle of training specificity because during the kick performance, martial athletes reach high angular velocity in the hip movement [6,17,18]. However, during the initial phases of the kick, the angular velocities are normally relatively low [6,17,18], and the lower limb segments acceleration occur at this moment, manifested at low contraction velocities during the start of the muscle torque production [7,19].

For the knee joint, the flexion and extension torques did not differentiate the two groups, and peak torque during the knee flexion or extension did not correlate with kinematical variables nor with the impact. These results show that, in expert athletes, the inter-individual differences in knee torque (see table 1) did not affect the competitive results. Presumably, this happened because the athletes in the present study did not significantly differ in their weekly training amount or in the total time spent training. Differently, Fong & Tsang [21] found significant correlations between the weekly specific taekwondo training and the knee flexion and isokinetic extension torques, but only at high speed (240°/s). Our results do not support the findings of Sbriccoli et al.[6] who found that for knee flexion elite karatekas produced higher torques than amateurs in all velocities varying from 30°/s to 400°/s.

Peak isokinetic torque during hip flexion and kicking time during the kick performance negatively correlated at both velocities of contraction. During the kick, shorter kicking time leaves the opponent less time to counterattack. Therefore, strong, and powerful hip flexors will generate kicks that are more effective. Such a finding is expected because, during the kicking phase, the dominant lower limb is off the ground and unconstrained, reducing the inertial resistance for the muscle action. It implies in facilitation for the action of the hip flexor muscles to produce velocity. The fast hip flexion movement during the kick and the isokinetic contraction with the fast contraction speed can be associated with the right-side of Hill muscle force-to-shortening velocity hyperbole, where the force is low. Additionally, during most of the kicking time, the hip flexes simultaneously to the knee, reducing the moment of inertia.

However, no significant correlations were found for the preparation time. Significant positive correlations were found between peak isokinetic torques during the hip flexion, extension and abduction, and the peak linear velocity of the foot during the kick. The correlations between joint torques and angular velocities were not usually significant, except for the hip extension at 60°/s and the knee flexion angular velocity during the kick.

A non-expected result was the negative correlation between the peak torque during hip adduction at 240°/s and the peak angular velocity of the foot during the kick. This can be due to the fact that hip adductor muscles activates during the preparation phase [27,32], and also flex the hip [33]. During the preparation phase, the hip is in a slightly extension position and the muscles might also participate to avoid a premature hip abduction while hip flexor torque is generated produced. Finally, the only torque values that significantly correlated with the impact magnitudes were produced during the hip flexion at 60 °/s.

The present study also shows a positive correlation between peak torque during the hip flexion at 240°/s and peak linear velocity of the foot during the kick. Sørensen et al.[7] have shown for the front kick that the hip flexors must counteract the hip extension reaction moment, caused by knee inertial torque during the knee angular acceleration towards extension. When the knee is extending, the hip angular velocity is relatively low [7], allowing larger hip flexion muscle forces, according to Hill’s hyperbole. The positive correlation between the hip flexor torque, at the slow isokinetic velocity, and the magnitude of the kick impact could also be expected because, during the impact, there is a high resistance of the movement, causing a sudden deceleration. This shifts the relationship force-velocity to the left of the Hill’s hyperbole.

The correlation found between the peak torque during the hip extension at 60°/s and the peak angular velocity during knee flexion (table 2) can possibly be explained by the biarticular function of the hamstrings, flexing the knee and extending the hip [33]. During the roundhouse kick, the biceps femoris activation occurs about 200 ms before significant production of muscle force, because of the electromechanical delay [26,27]. For this reason, the hamstrings’ activation and the start of the knee flexion action occur before the kicking phase, simultaneously with the hip extension of the preparation phase [26,27]. The correlation between the hip extensor torque at slow isokinetic speed and the foot linear velocity can be explained by the proximal-distal momentum transmission [7,34]. The action of hip extensor muscles, in the correct time can potentiate the proximal-distal momentum transmission as demonstrated in our previous studies [27,32].

A significant correlation was found between the peak torque during the hip abduction at 240°/s and the peak lineal foot velocity. Hip abduction torque accelerates the thigh and, consequently, the foot segment. Before the impact, the knee extends and suddenly increases the moment of inertia of the entire leg relative to the hip. At this moment, the hip abducts, and the increased moment of inertia acts toward reducing the angular acceleration. Therefore, stronger abductors will probably provide more speed to the foot during the kick and explaining the significant correlation. All the correlations obtained in this study had low to moderate effect sizes, then, the most considerable portion of the kick performance in taekwondo may be associated with other parameters, such as the technique and coordination. Some studies confirm the importance of coordination for the kick speed and impact intensity, by demonstrating that the kick performance is influenced by factors such as intra-limb [16] and inter-joint coordination [10], momentum (and kinetic energy) proximal-to-distal transmission [7,9,34], muscle cocontraction [4,6,17,26,27] and using the stretch-shortening cycle [15,34].

The present study also identify three significant discriminant equations (table 3) for the expertise level based on: a) peak torques during isokinetic muscle contraction; b) kinematic variables during kicking performance and; c) a combination of both. The last equation (Equation “D_3_”) supports the third hypothesis of the study, such that a discriminant equation combining kinematic and torque variables would be a strong predictor of the ranking performance. For this equation, based on hip flexion and extension torques at 240°/s and on the knee angular velocity, we found a high level of statistical performance for all parameters of the group prediction (elite or sub-elite). Namely, accuracy (“hit-ratio”), sensitivity and specificity obtained a success rate of 85.7% for the expertise level. The isokinetic torque variables represent the muscle force and power capacity to generate torque in a simple (monoarticular) task, relatively independent of the motor coordination and the technical specific skill [6,34]. The angular velocity of the knee extension during a kick represents the opposite, i.e., the capacity to use muscle force, influenced by technical skills [10,15,16,34], to generate a sport-specific performance. Such a neuro-motor task should include the ability for taking advantage of the proximal-to-distal momentum transmission [7,9,34]. For example, from the discriminant function D_2_, the angular velocity borderline to discriminate elite from sub-elite athletes is a good predictor to categorize the athletes in their respective group. That is, those scoring above and below than 1504 deg/s can be categorized as elite or sub-elite, respectively.

As limitations of this study we would like to highlight that the isokinetic hip torque can be associated with the specific performance of TKD. Future longitudinal studies are necessary to confirm the effect of a designed training for improving muscle torques of selected joints on the specific performance of TKD movements. Also, inverse dynamics to calculate the torques and transmitted momentum from proximal to distal segments during the kick was not performed, which could have enriched the discussion of the results.

## Conclusion

Isokinetic hip flexion and extension torques are associated with specific TKD performance. Such torques are important for both the differentiation of the competitive ranking of athletes and for the correlations with the kinematic performance and impact of the kicks. The capacity of muscle force production at slow speed (60°/s), and not only at high contraction speed (240°/s), is relevant for the taekwondo performance. Slow speed torques significantly correlated with critical parameters of the kick performance, including the linear foot velocity, impact magnitude, and kick time. Additionally, the combination of isokinetic torque production (hip sagittal torque at 240°/s) with a sport-specific kinematic performance parameter (angular velocity of knee extension), using a linear discriminant function, yielded a strong predictor of the athletes’ expertise level.

### Practical implications

a. Experienced taekwondo athletes that aim to improve their competitive level and training efficiency should strengthen the muscles that control the hip joint movements (flexion, extension, and abduction).
b. This strength training should be performed both at low loads with high speed and at high loads with relatively low speed. Through this, we can speculate that training the hip muscles strength with overload (elastic tubes, free weights, pulley exercise, etc.) has the potential to improve the kick impact, more than to train other muscle groups.
c. The most considerable portion of the kick performance in taekwondo may be associated with other parameters, such as the technique and coordination.

## Acknowledgment

I thank the masters Alan do Carmo and Carmen Carolina for kindly granting their best athletes to carry out the research.

## Supplementary Material

### Supplementary Media Files

**Video 1 -** Demonstration of the data collection and the rigid body diagram of one athlete performing a Bandal Chagui kick followed by a kinematic analysis resulted from a number of Bandal Chagui kicks performed by another athlete of the study. Figshare, Uploaded in: September 04, 2019. Available at: https://doi.org/10.6084/m9.figshare.9698741.v2.

### Supplementary Data

**S1_Data.sav –** Data for statistical analysis in SPSS spreadsheet.

**Legend: PIT**, Peak of Isokinetic Torque; **AVK**, Angular Velocity during the kick movement; **PT**, Preparation Time, the impulse phase of the kick; **KT**, Kick Time, the aerial phase of the kick; **LVF**, Linear Foot Velocity; **LVK**, Linear Knee Velocity; **LVP**, Linear Pelvis Velocity; **KnFlx**, Knee Flexion; **KnExt**, Knee Extension; **HipFlx**, Hip Flexion; **HipExt**, Hip Extension; **HipAdd**, Hip Adduction; **HipAbd**, Hip Abduction; **WTU**, World Taekwondo Federation (WT) patented Unity (U) of Power Impact. **60**. Degrees for second on isokinetic evaluation and **240**, 240 degrees for second on isokinetic evaluation.

